# A Pipeline for Faecal Host DNA Analysis by Absolute Quantification of LINE-1 and Mitochondrial Genomic Elements Using ddPCR

**DOI:** 10.1101/522862

**Authors:** Kuang He, Hideaki Fujiwara, Cynthia Zajac, Erin Sandford, Pavan Reddy, Sung Won Choi, Muneesh Tewari

**Affiliations:** Division of Hematology and Oncology, Department of Internal Medicine, Comprehensive Cancer Center, University of Michigan, Ann Arbor, Michigan, 48109, USA; Department of Pediatrics & Communicable Diseases, University of Michigan, Ann Arbor, Michigan, 48109, USA; Center for Computational Medicine and Bioinformatics, University of Michigan, Ann Arbor, Michigan, 48109, USA; Department of Biomedical Engineering, University of Michigan, Ann Arbor, Michigan, 48109, USA; Biointerfaces Institute, University of Michigan, Ann Arbor, Michigan, 48109, USA

## Abstract

Stool contains DNA shed from cells of the gastrointestinal (GI) tract and has great potential as a bio-specimen for non-invasive, nucleic acid-based detection of GI diseases. Whereas methods for studying faecal microbiome DNA are plentiful, there is a lack of well-characterised procedures for stabilisation, isolation, and quantitative analysis of faecal host DNA. We report an optimised pipeline for faecal host DNA analysis from the point-of-collection to droplet digital PCR (ddPCR) absolute quantification of host-specific gene targets. We evaluated multiple methods for preservation and isolation of host DNA from stool to identify the highest performing methods. To quantify host DNA even if present in partially degraded form, we developed sensitive, human-specific short-amplicon ddPCR assays targeting repetitive nuclear genomic elements (LINE-1) and mitochondrial genes. We validated the ability of these optimised methods to perform absolute quantification of host DNA in 200 stool DNA extracts from samples that were serially collected from three healthy individuals and three hospitalised patients. These specimens allowed assessment of host DNA day-to-day variability in stool specimens with widely varying physical characteristics (i.e., Bristol scores). We further extended this approach to mouse stool analysis, to enable faecal host DNA studies in animal disease models as well.

## Introduction

Analysis of DNA in stool has attracted great interest, which has largely focused on the gut microbiota and its relationship to health and disease. Aside from microbes, stool also contains exfoliated cells from the lining of the gastrointestinal (GI) tract ^1^. Given that both genetic ^2,3^ and epigenetic ^4^ changes in DNA of somatic cells underlie many diseases, stool DNA tests offer great opportunities for non-invasive sampling and study of the GI tract in health and disease, as shown by commercial success of Cologuard (Exact Sciences, Inc.), a stool tumour DNA-based test for early detection of colorectal cancer. However, unlike the microbiome field where methods for preservation, isolation, and quantitative analysis of stool microbial DNA are well-established, comparable well-characterised methods for the study of host DNA in stool are lacking in the public domain. Challenges to the successful analysis of host DNA in stool include the lack of:

- <u>Sample preservation at the point of collection for host DNA stabilisation</u>: for the microbiome, immediate freezing of stool samples or storage in specific preservative solutions before DNA extraction significantly improves the stability of microbial community compositions compared to no preservation. For human DNA preservation in stool, there have been a few studies that assessed preservation in EDTA-based buffers and commercial solutions ^6–8^. However, it was unclear whether DNA stabilisation solutions reported earlier are effective in preserving a range of DNA fragment lengths, including short fragments of host DNA that may be derived from normal apoptotic colonocytes or neoplastic cells (i.e. ~100 bp) ^9^.
- <u>High efficiency host DNA extraction</u>: most commercial methods for stool DNA extraction are optimised for long microbial genomic DNA and not for host DNA, which also includes the shorter host DNA fragments expected from apoptotic epithelial cells shed into the stool. In addition, some of the earlier work has been done with proprietary commercial reagents that are not easily accessible for research use ^8^. Thus, there is a need for detailed studies of the efficiency of various stool DNA isolation methods for recovering host DNA, in the public domain.
- <u>Assays for absolute quantification of host DNA targets</u>: the low abundance of host DNA (typically <1% total stool DNA ^10,11^) and the presence of PCR inhibitors of dietary and metabolic origin can present challenges for high sensitivity absolute quantification of host DNA using traditional quantitative PCR (qPCR) methods.

To address these challenges, we studied three preservation solutions for human DNA stabilisation during stool collection and transportation, evaluated three commercially available DNA isolation kits for their ability to recover DNA without size bias, and developed sensitive nuclear and mitochondrial DNA element assays to human DNA in stool using droplet digital PCR (ddPCR). ddPCR enables single DNA molecule detection by partitioning PCR reactions into many thousands of oil-capsulated nanolitre-sized droplets and performing PCR amplification in individual droplets. ddPCR is well-suited for host DNA quantification, as it is an absolute quantification technique that more robust to PCR inhibitors than qPCR, and offers greater precision and improved day-to-day reproducibility than qPCR without requiring a standard curve ^13, 14^. Here we report an optimised pipeline using 0.5 M EDTA (pH 8) for stool preservation, specialised reagents for DNA extraction (Norgen Biotek Corp.), and ddPCR of LINE-1 and mitochondrial DNA targets to perform absolute quantification of host DNA in stool. We report data from not only healthy individuals, but also hospitalised patients (i.e., recipients of allogeneic hematopoietic cell transplantation (HCT)) who often experience GI disturbances (e.g., diarrhoea) that result in stools of a range of physical characteristics (i.e., Bristol scores). Finally, we developed and validated assays for host DNA analysis in stools from mice, to enable study of host DNA in stool samples from a commonly used animal model system.

## Results

### Design and Optimisation of ddPCR Assays for High Specificity Detection of Human DNA in Stool

Human DNA is present in relatively low abundance in stool ^10,11^ and is expected to have varying lengths due to naturally occurring apoptosis of colonic epithelial cells ^15^ and potential degradation by nucleases ^16,17^ present in stool. Therefore, when selecting human gene targets and designing ddPCR primers for our assays, we considered: i) the gene targets should ideally exist as a large number of copies per cell for enhanced sensitivity; ii) the primers need to be highly specific for human DNA relative to microbial, plant, or animal genomes that may be present in stool; and iii) the amplicons need to be as short as possible to allow for efficient capture of even partially degraded DNA. Therefore, we focused on two types of targets that are present at multiple copies per cell: repetitive sequences in the nuclear genome and mitochondrial genes. Long interspersed nuclear elements (LINEs), including LINE-1 repeats, are transposable elements that comprise 17% of the human genome ^18^. LINE-1 repeats in plasma have been used to quantify the human tumour xenograft load in mice ^19^. We postulated that the extremely high copies per genome of LINE-1 elements would substantially increase the likelihood of detecting human DNA using these targets, even in a high background of microbial and dietary DNA. Mitochondrial (mt) DNA is another desirable target because mitochondrial DNA sequences can be species-specific, and could therefore provide a human-specific DNA target. In fact, mtDNA markers have been used to track faecal contamination in water environments ^20^. Furthermore, there are tens to thousands of mitochondria per human cell, depending on the cell type ^21^, which could enhance sensitivity. We therefore selected two LINE-1 sequences, 55-bp and 60-bp long, respectively, as well as two mitochondrial gene sequences, 83-bp (located in the NADH dehydrogenase subunit 5 gene, abbreviated as ND5) and 77-bp (located in the cytochrome c oxidase subunit II gene, abbreviated as CO2) long, respectively, as targets for initial ddPCR assay development.

We first determined optimal annealing/extension temperatures for the four primer sets (described in Materials and Sequences) using extracted human stool DNA as a template in ddPCR reactions. For the LINE-1 primer sets, we observed decreasing amplification as annealing/extension temperatures were increased from 53 to 61 °C (Fig. 1a). We selected 60 °C as a conservative temperature to minimize false positive signals. For the mt primers, on the other hand, absolute copy number (ACN) quantifications remained relatively constant between 53 - 61 °C (Fig. 1a). We decided to use the same annealing/extension temperature of 60 °C for ddPCR for both primer sets so these two assays can be run together on the same plate. At 60 °C, LINE-1 assays yielded 6-20 times higher ACN than mt assays, with the LINE-1 60-bp amplicon demonstrating the highest ACN of all (Fig. 1a), suggesting that LINE-1 assays may be more sensitive for low DNA input samples than the mt assays.

**Figure 1.**
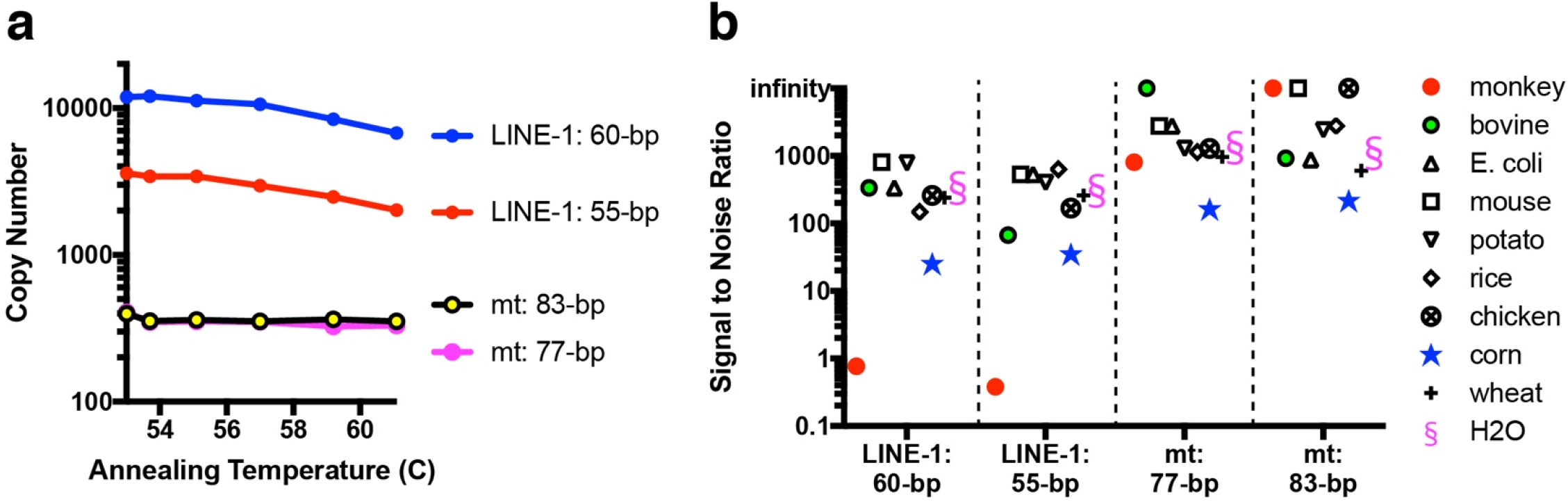
Optimisation of PCR assay conditions and specificity assessment for the faecal human DNA ddPCR assays. (a) Annealing temperature optimisation and (b) DNA specificity evaluation of two sets of ddPCR primers targeting human LINE-1 elements and two sets of ddPCR primers targeting human mt genes. (a) 80 pg of total stool DNA extracted from a human stool sample was used as template in all four gradient PCR reactions. (b) S/N ratio represents ACN obtained from 80 pg of total stool DNA extracted from a human stool sample, compared to ACN from 80 pg of purified gDNA from various animal, plant, and bacterial sources, as well as water. The calculation of signal to noise ratios is defined in the text.

In order to assess specificity of the assays for human DNA, relative to other types of genomes that may be present in stool, we ran the four assays on 80 pg extracted human stool DNA as well as 80 pg of genomic DNA (gDNA) purified from a range of animal, plant, and bacterial sources. For each non-human genome, we determined a signal to noise (S/N) ratio calculated as ACN measured in human stool DNA divided by ACN measured from the non-human gDNA. As shown in Fig. 1b, for LINE-1 assays, aside from monkey gDNA (where cross-reactivity was expected given sequence conservation), the assays showed human-specificity in excess of 100-fold across most genomes, and even the lowest S/N ratio, seen with one of the assays on corn gDNA, showed in excess of 25-fold specificity for human gDNA. The mitochondrial DNA assays were even more human-specific, yielding S/N ratios of >100 in all cases and >1000 for some of the non-human genomes (Fig. 1b). Whereas the mtDNA assays show higher specificity, the LINE-1 assays may be more sensitive, because of the potentially higher number of copies per cell of LINE-1 DNA targets. Based on the results in Fig. 1b, we selected the 60-bp LINE-1 amplicon and 83-bp mt amplicon for subsequent assay development and characterisation.

### ddPCR Linearity, Accuracy, and Reproducibility

To test the linearity, accuracy, and reproducibility of the selected LINE-1 and mt assays, we used purified human gDNA and synthetic gBlock fragments as templates to generate triplicate standard curves for each assay on three different days. Since the manufacturer (Bio-Rad, Inc.) recommends DNA fragmentation of genomic DNA with restriction enzymes for ACN analysis, we digested purified human gDNA with the restriction enzyme *Hae*III, which yields a theoretical average fragment size of 347 bp^22^.

Using synthetic gBlocks as templates, we found that our assays were linear in the range from 193 to 120,440 copies of input DNA target molecules, with a median % yield (i.e., (# of copies measured/# of copies expected)*100) of 50.6% for the LINE-1 assay and 44.1% for the mt assay (Fig. 2a-b **and Supplementary Table S1**). It is worth noting that the gBlock working solutions used for this analysis were stored as highly dilute solutions (10 pM) to reduce the risk of cross contamination in the lab. This may have contributed to reduced % yield, via DNA adsorption to the plastic surface of the storage tubes that can occur with dilute solutions. Using digested gDNA as templates, we found that our LINE-1 assay is linear in a range from 36 to 22,247 *in silico* predicted copies input, with a median % yield of 395% (Fig. 2c **and Supplementary Table S1**). Our experimental results indicate that the *in silico* predictions are an underestimate. The expected (i.e., theoretical) yield value for the LINE-1 PCR used was ~5,800 copies per genome, as determined using an *in silico* PCR simulation tool available on the UCSC Genome Browser^23^. However, with highly abundant repeat elements containing somewhat degenerate sequences throughout the genome, it is difficult to know the precise accuracy of *in silico* tools. Regardless, the data indicate that the assay detection limit lies well below the amount of DNA corresponding to a single cell (i.e., less than one genome equivalent (GE)). The standard curves demonstrated excellent linearity for both the LINE-1 primer set (Fig. 2a, 2c) and mt primer set (Fig. 2b).

**Figure 2.**
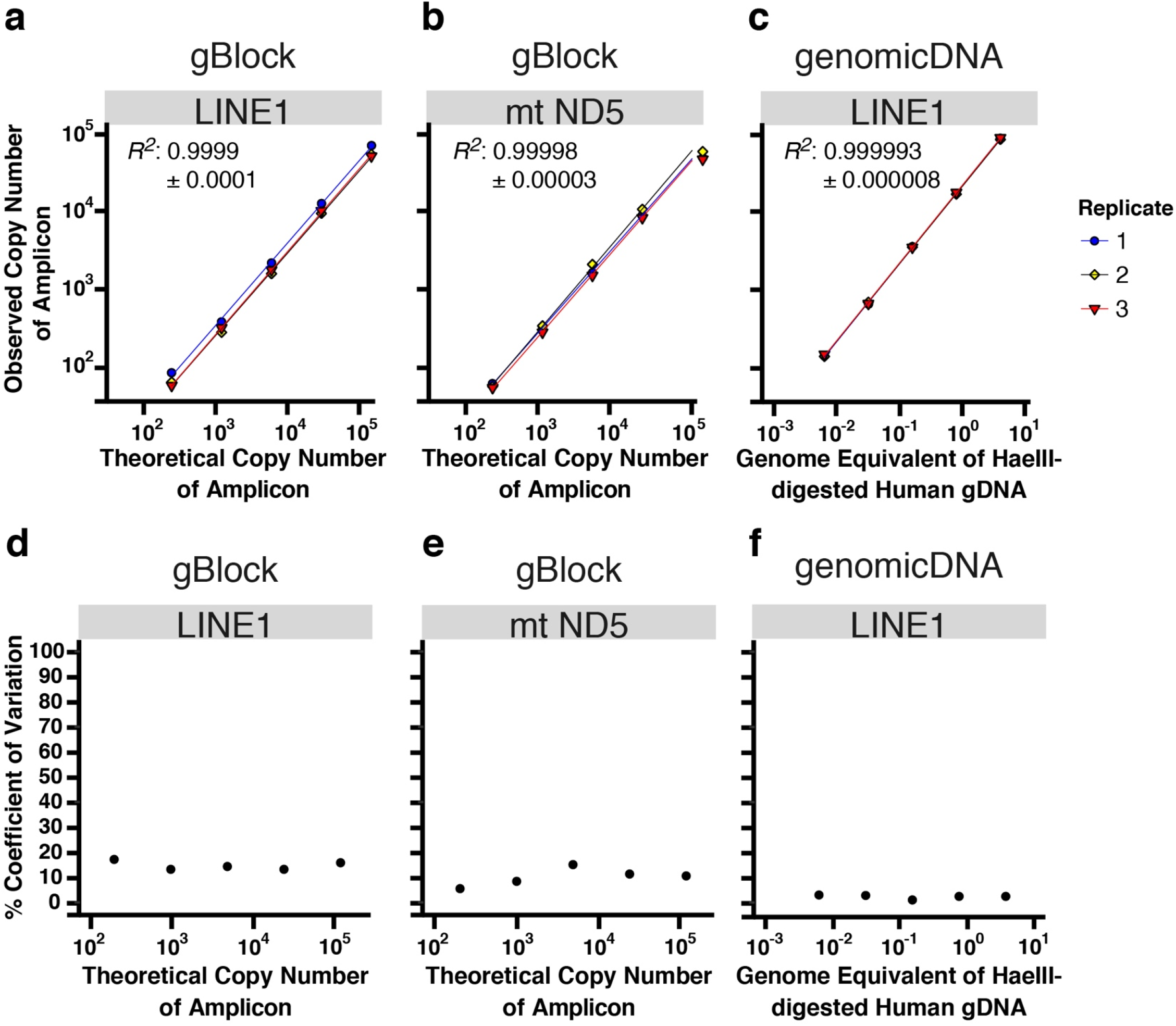
Analytical method validation of the ddPCR assays. Linearity (a-c) and reproducibility (d-f) of the LINE-1 and mt assays were assessed in triplicate using synthetic gBlocks and digested human gDNA as templates. Data are fitted to linear regression models. Theoretical copy number represents the expected number of molecules based upon the known concentration of the synthetic gBlock solution.

To assess day-to-day reproducibility, we calculated percentage coefficients of variation (% CV) for the triplicate standard curves performed on three different days as shown in Fig. 2d-f. The % CVs were 5-18% when concentrated synthetic gBlocks were used as templates for LINE-1 (Fig. 2d) and mt assays (Fig. 2e), and <3% at each of the five tested concentrations of human gDNA for the LINE-1 assay (Fig. 2f). We attribute the higher % CV seen with the gBlocks to the fact that these are prepared as high-concentration stock solutions requiring more serial dilution steps than the gDNA templates, which likely incurred greater pipetting variations.

### Evaluating Stool DNA Purification Methods for High Efficiency Recovery of Host DNA

Maximum DNA recovery during DNA purification is critical for accurate DNA quantification. As it is highly likely that *in vivo* (i.e., inside intestines) as well as *ex vivo* (i.e., during specimen collection, transportation, and storage) degradation and fragmentation of host DNA might occur in stool, we searched for DNA isolation methods likely to recover short and long DNA fragments equally well. We tested three commercial stool DNA purification kits (the Zymo Quick-DNA faecal/Soil Microbe Miniprep Kit, the Norgen Stool DNA Isolation Kit, and the Qiagen QIAamp DNA Stool Mini Kit (human DNA protocol)) for recovery of a DNA ladder (Invitrogen 1kb Plus), ranging from 100 to 15,000 bp. By loading DNA ladder either <u>d</u>irectly (D) onto the TapeStation, or after one of the three purification protocols (P) (Fig. 3a), we visualised DNA losses by looking at the differences in band intensity of the DNA ladder components before and after <u>p</u>urification (Fig. 3b). We chose 10 g and 2 g as total input DNA amounts for purification because these are similar in range to the amount of stool DNA we expect to be present in the aliquots of stool specimens that we would typically analyze, taking into account historical stool DNA abundance data from prior work ^24^.

**Figure 3.**
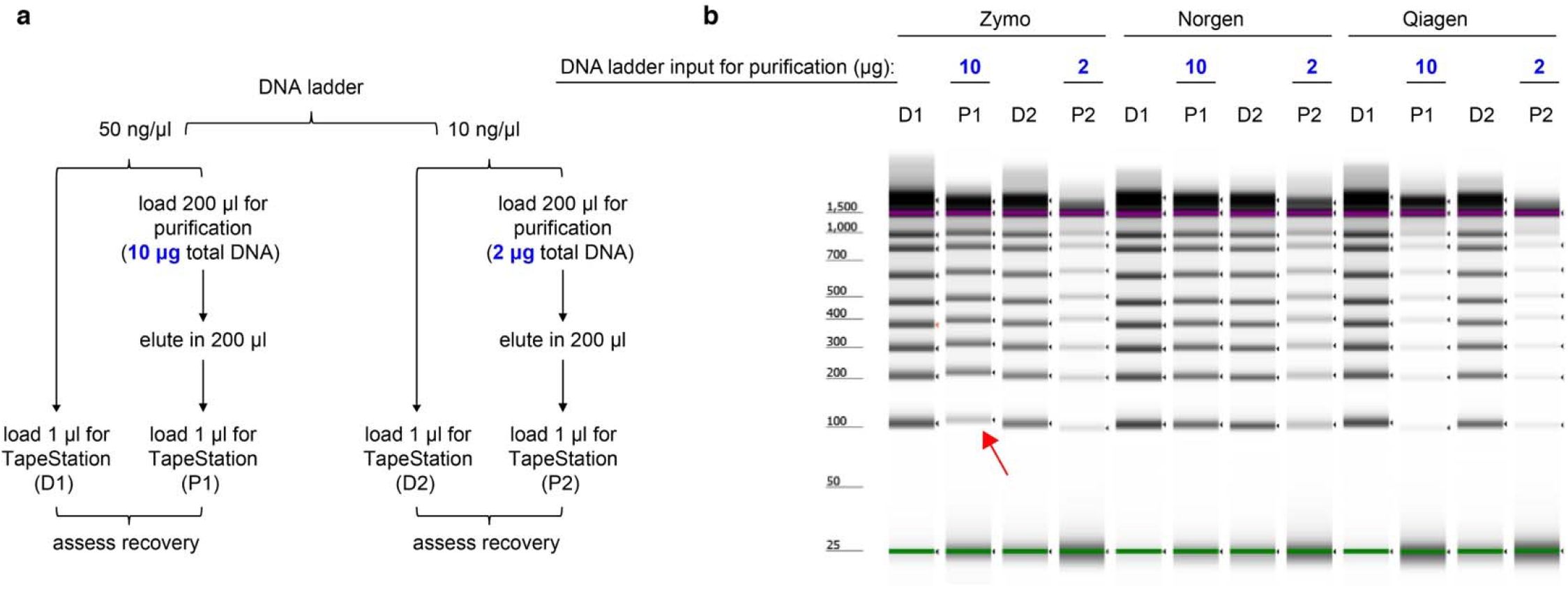
Comparison of DNA recovery of a ladder consisting of fragments as short as 100 bp using the Zymo Quick-DNA faecal/Soil Microbe Miniprep Kit (Zymo), the Norgen Stool DNA Isolation Kit (Norgen), and the QIAamp DNA Stool Mini Kit (Qiagen). (a) For direct visualization of DNA recovery from the three different purification processes, DNA ladder was either loaded directly onto TapeStation (D) or was subjected to one of the three purification protocols before being loaded onto TapeStation (P). Please note that band intensities between lanes D1 and D2 are not directly comparable because the higher input into D1 lanes may be beyond the linear range of the assay. In a 100% recovery scenario, the D and P lanes should contain the same amount of DNA. (b) A representative image of gel electrophoresis on TapeStation. The size marker lane was cropped out for clarity (the uncropped, full-length gel is presented in **Supplementary Fig. S1**). The purple and green lines in each lane represent internal upper and lower markers, respectively, for sizing and alignment. The red arrow denotes low recovery of the 100 bp band from the Zymo-extracted ladder samples.

At the 10 μg DNA input level, the Norgen reagents showed the most uniform and highest recovery of DNA fragments, Zymo was comparable in recovery across the DNA ladder range except it showed diminished recovery of the 100 bp fragment (see red arrow in Fig. 3b), and the Qiagen kit showed the lowest recovery of DNA across the entire size range. At the 2 μg DNA input level, the Norgen reagents showed the highest recovery (Fig. 3b). Based on these results, we selected the Norgen DNA isolation method for further analyses because of its high efficiency, uniform DNA recovery of the full range of sizes examined.

We then assessed DNA isolation efficiency of the Norgen kit using healthy donor human stools as a background and *Hae*III-digested human gDNA as spike-ins. Human stools were lysed using Norgen reagents, centrifuged, and supernatants aliquoted in replicates. Serially diluted human gDNA or a buffer control was mixed with the lysate aliquots and carried through the rest of the isolation protocol per the manufacturer’s instructions. The LINE-1 assay was used to quantify the ACN of the 60-bp amplicon in each sample, with and without the spike-ins. The percentage of DNA spike-in recovery, R, was calculated as:

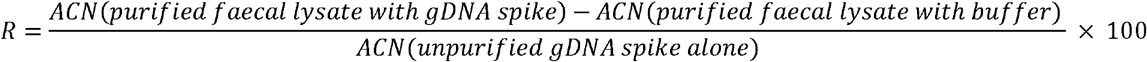

As shown in Fig. 4, in the background of total stool DNA, 800 ng, 80 ng, and 8 ng human gDNA spike-ins corresponding to 232,000 GE, 23,200 GE, and 2,320 GE resulted in an average of 57% ± 5%, 60% ± 11%, and 75% ± 18% recovery, respectively, through the DNA isolation process. These values give us confidence that the majority of human DNA in stool can be recovered for downstream analysis.

**Figure 4.**
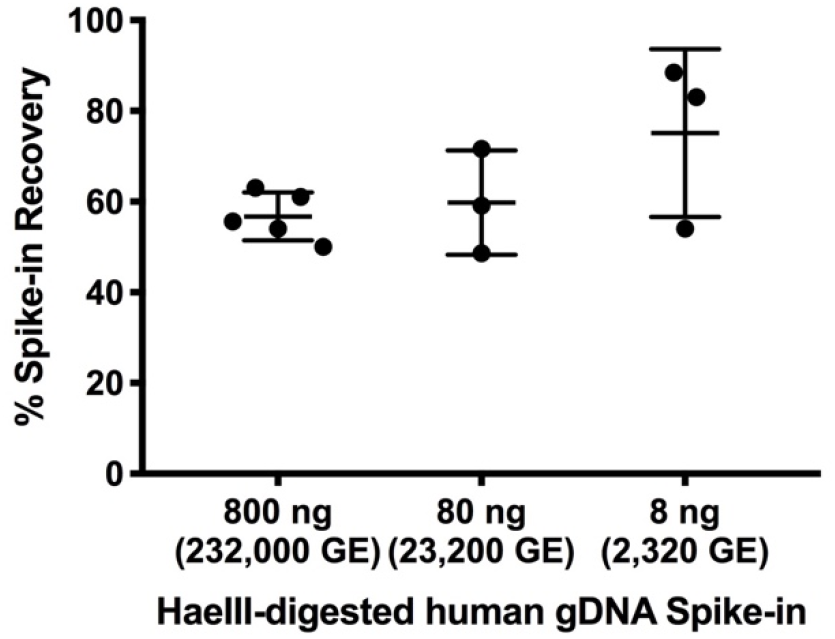
DNA isolation efficiency using the Norgen Stool DNA Isolation Kit. Across all samples (ranging from 800 to 8 ng of digested human gDNA spiked into stool slurries and carried through DNA isolation), the recovery of the spike-ins averaged 62%. Error bars represent standard deviations.

### Selecting a Host DNA Preservative Buffer for Stool Collection

Considering that stool transport after collection most conveniently occurs at room temperature, we sought to evaluate preservative solutions for stool host DNA stabilisation. We aimed to assess time-dependent DNA degradation of homogenised, buffer-preserved stool at room temperature to simulate a typical specimen transport temperature. Buffers selected for this study are: (i) a proprietary buffer OMNIgene (referred to as OMNI hereafter), which comes with the OMNIgene Gut Kit and has been optimised for microbial DNA ^25^, (ii) a buffer referred to as TEN2 which contains Tris, EDTA, and NaCl, which represent core ingredients of a previously described stool DNA preservative solution ^26,27^ and (iii) a simple solution of 0.5 M EDTA at pH 8.0 (referred to as EDTA hereafter) designed to inactivate DNases by chelating divalent cations.

We collected stools from two healthy individuals (D-159x and D-145x), who scooped freshly defecated stools into collection devices containing OMNI, TEN2, or EDTA solutions. Stool specimens were brought to the laboratory within an hour. Stools were subsequently weighed, the buffer volume adjusted (for TEN2 only, see Materials and Methods), homogenised, and then aliquoted into five portions. Each aliquot was frozen at −80 °C after incubation at room temperature (22 °C) for one of five different time durations (0, 4, 24, 72, and 96 hours). See Fig. 5a for a schematic diagram of the workflow. We then extracted faecal DNA from each of the time point samples using Norgen reagents and used ddPCR to measure LINE-1, mt, and bacterial DNA targets as per methods described above.

**Figure 5.**
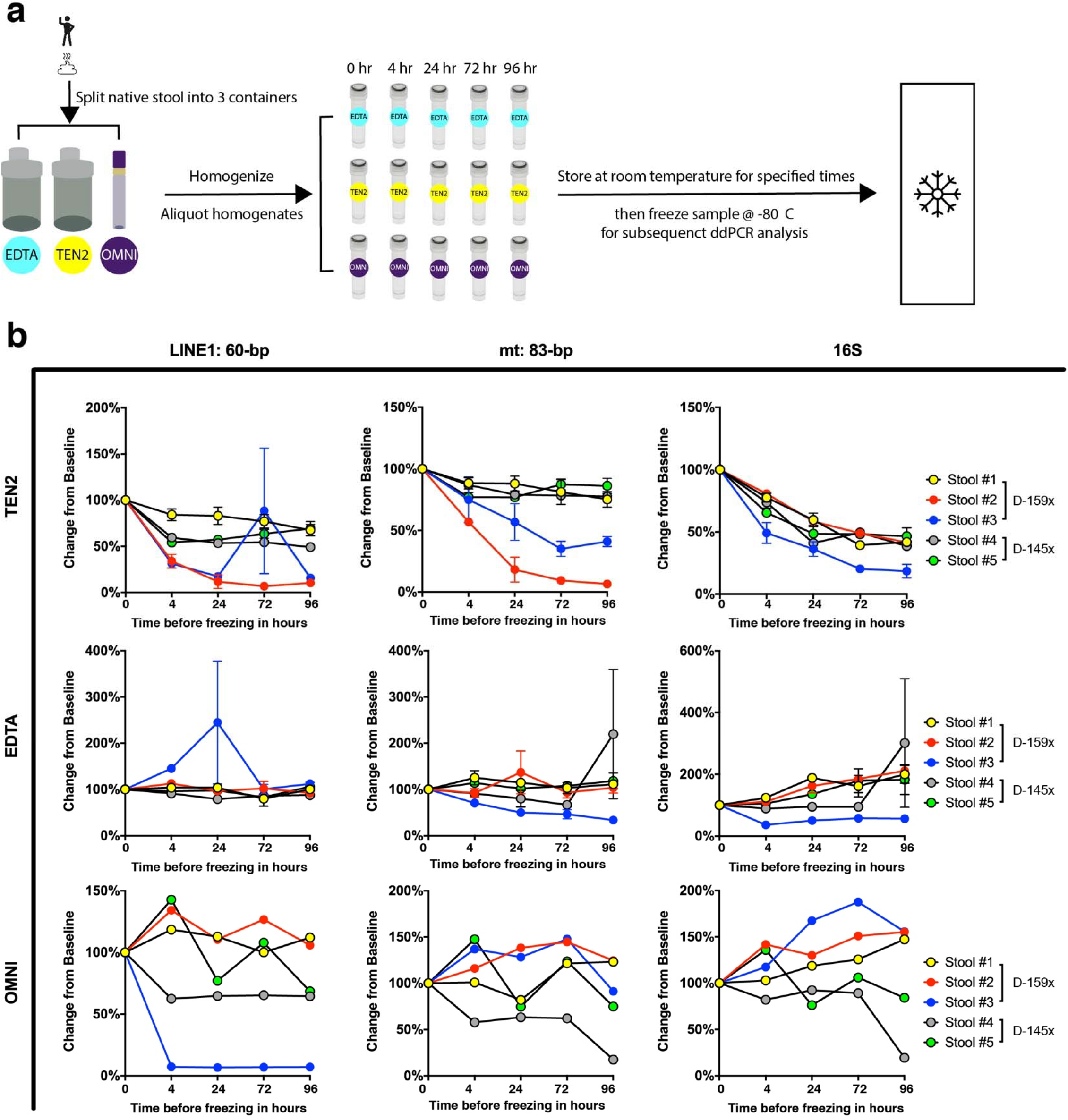
Evaluation of three different stool preservative solutions for host DNA stabilisation at room temperature. (a) Schematic diagram of stool processing for the stool DNA stability experiment and (b) Effectiveness of different stool preservation solutions on endogenous DNA stability. Change in ACN per μl stool DNA extract from baseline (time 0) of LINE-1 (left), mt (ND5) (middle), and 16S (right) DNA are plotted for varying incubation times of stool specimens at room temperature. TEN2- and EDTA-preserved stools were DNA extracted and analyzed in duplicate; the error bars represent standard errors from ddPCR of two replicate extractions for each time point. The OMNI-preserved stool provided enough material only for one extraction per time point and therefore error bars are not provided.

As seen in Fig. 5b (relative change in ACN per μl extract from baseline, plotted against storage time) and **Supplementary Fig. S2** (ACN per μl extract plotted against storage time), there are differences in the stability of DNA in stools preserved in different buffers over time. In TEN2, ACN of both human and microbial DNA targets decreased over the course of four days storage at room temperature, indicating progressive DNA degradation. In EDTA, we found that the ACN of human genes tended to stay constant over time, and that of bacterial genes rose slightly over the four days, indicating modest growth of faecal bacteria. And in some samples preserved in OMNI, levels of host DNA decreased whereas those of bacterial DNA increased over the incubation period. Based on these results, we selected EDTA as the stool preservation solution for subsequent experiments.

### Quantifying Host DNA in Serially Collected Stool Specimens from Healthy Individuals

To demonstrate the feasibility of using the optimised procedures developed here for assaying human DNA in stool, we applied them to analyze stool specimens in duplicate from three healthy control individuals (D-145x, D-165x, and D-166x), collected on multiple days (31 samples and 62 DNA isolations in total). These individuals had no known GI disorders. Using our DNA isolation protocol, the median stool input per ~200 μl stool homogenate for DNA extractions was 49.5 mg (Fig. 6a), and the median recovered total DNA per extraction was 707 ng (Fig. 6b). Figure 6c-d show the ACN per μl extract and normalized as per mg stool, respectively, of human-specific LINE-1 targets recovered from these samples. Figure 6e-f show the ACN per μl extract and normalized as per mg stool, respectively, of human-specific mtDNA targets recovered from these samples. Both LINE-1 and mtDNA targets were detected in all samples, with measured mtDNA levels being 9-fold lower than LINE-1, on average. Longitudinally, we observed a several-fold intra-individual day-to-day variation in human DNA levels in stool in the healthy individuals. This was not affected by normalisation to stool mass (i.e., ACN per mg stool input) (Fig. 6c-f), indicating that the observed day-to-day variation represents true biological variation.

**Figure 6.**
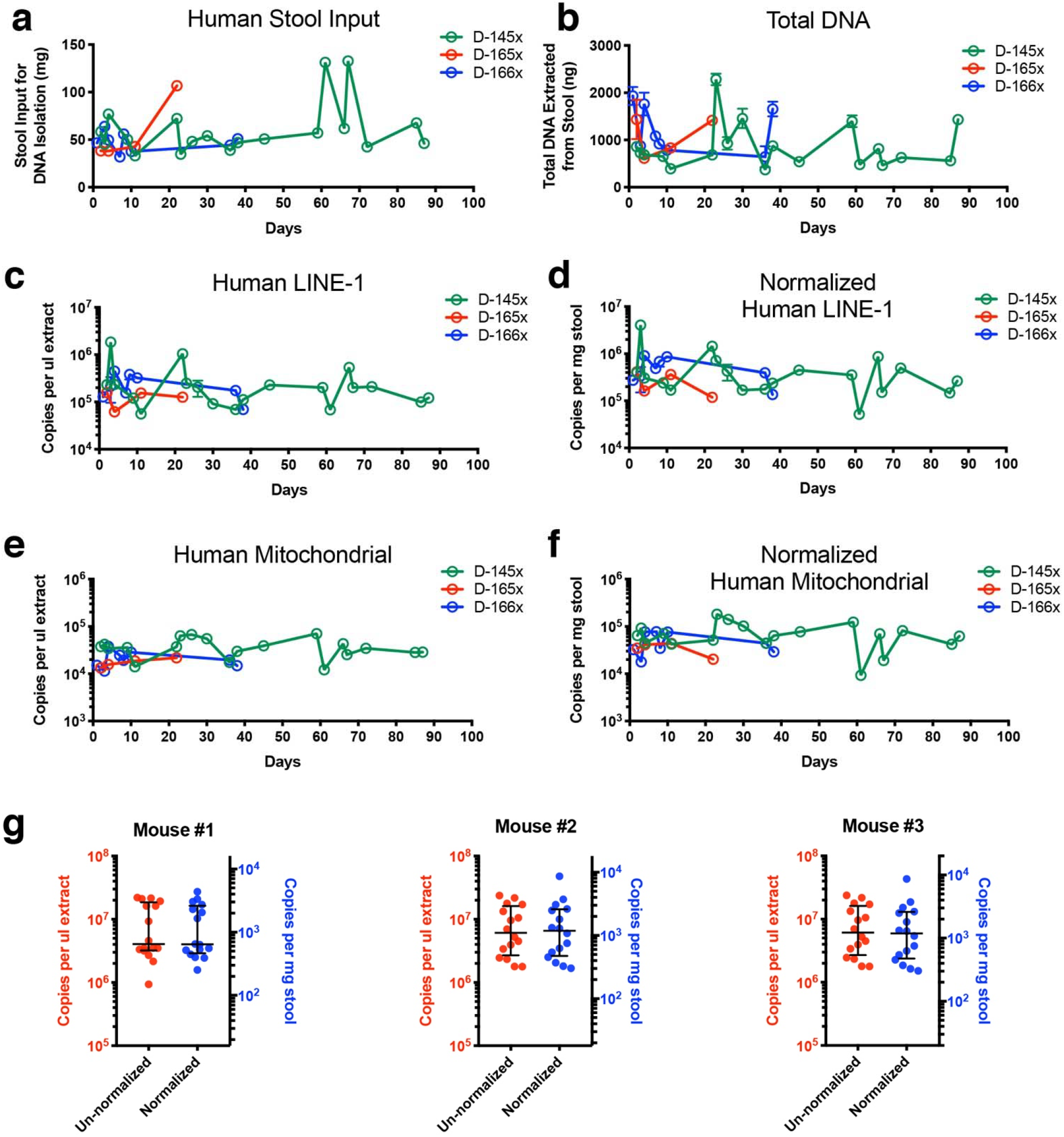
Quantification of host DNA from healthy human and murine stools using ddPCR. (a) Plot showing the variation in amount of stool input used for DNA purification for each of the longitudinally collected stool specimens from three healthy humans (D-145x, D-165x, and D-166x). (b) total DNA isolated from the amounts of stool shown in (a). The error bars represent the range of total DNA isolated from two side-by-side purifications per specimen. (c) Human DNA quantified from total fecal DNA extracts using the human LINE-1 (60-bp amplicon) assay, expressed in copies per μl DNA extract. (d) Human DNA quantified from total fecal DNA extracts using the human LINE-1 (60-bp amplicon) assay, normalized by expressing as copies per mg stool. (e) Human DNA quantified from total fecal DNA extracts using the human mt (83-bp amplicon in the ND5 gene) assay, expressed in copies per μl DNA extract. (f) Human DNA quantified from total fecal DNA extracts using the human mt (83-bp amplicon in the ND5 gene) assay, normalized by expressing as copies per mg stool. Error bars in (c-f) represent range of copy numbers from single ddPCR reactions on the duplicate DNA extraction. (g) Quantification of mouse DNA in stools collected from three healthy mice using the mouse LINE-1 (58-bp amplicon) assay. LINE-1 levels are plotted as copies per μl stool DNA extract and copies per mg stool for each mouse. The horizontal lines represent the median and interquartile range.

Based on our success in detecting host DNA in human stool, we next applied the approach to mouse stool DNA quantification. Two sets of primers targeting LINE-1 elements in the mouse genome (mLINE-1) were designed. These mLINE-1 primers target 58-bp and 62-bp amplicons, respectively. We used 0.2 pg HaeIII-digested purified mouse gDNA as templates in a gradient PCR experiment to determine the optimal annealing and extension temperatures for the primers. Similar to the human LINE-1 primers, the mLINE-1 primers showed inverse relationships between copy number yields and annealing and extension temperatures (**Supplementary Fig. S3a**). To minimise non-specific amplification and ensure proper primer-template annealing, we chose 59 °C for annealing and extension during ddPCR, a temperature at which both primer sets exhibit high copy number yields (~10,000 copies/GE for the 58-bp amplicon and ~16,300/GE for the 62-bp amplicon).

To test the specificity of the two mLINE-1 primer sets, we included DNA isolated from mouse chow in addition to the genomes we tested for the human primer sets. Signal to noise ratios for each non-mouse DNA source were calculated as ACN per pg DNA from the mouse stool sample, divided by ACN per pg DNA from each of the non-mouse genomic DNA. As shown in **Supplementary Fig. S3b**, both primer sets exhibit signal to noise ratios of >1000 against all non-mouse DNA. Therefore, both mLINE-1 assays are highly specific to mice.

Stool pellets were collected from three healthy BALB/c mice twice a week for five weeks. We collected fresh pellets directly from the anus of the mice into tubes containing EDTA. Using the primers targeting the 58-bp mLINE-1, we plotted the un-normalised stool mLINE-1 ACN for each mouse (Fig. 6g, **left y axes**) as well as ACN normalised to mg of input stools (Fig. 6g, **right y axes**) and observed a several-fold day-to-day variation in host DNA levels, similar to the variations seen in healthy human LINE-1 measurements (Fig. 6c-d).

### Quantifying Host DNA in Stool Specimens from Hospitalised Patients

In contrast to stool from healthy individuals, stool from hospitalised patients experiencing acute and chronic GI tissue damage or inflammation can vary greatly with respect to physical consistency (e.g., ranging from firm and dense to entirely liquid), the presence of potential PCR inhibitors^28^, and microbiome mass^11^ and composition^29^. All of these factors could collectively alter host DNA content and accessibility for analysis in the stool. To determine whether our optimised methods could be used to measure host DNA levels in stool from hospitalised patients, we used our methods to quantify human DNA in three allogeneic hematopoietic cell transplantation (HCT) patients (P07, P12, and P13) from duplicate DNA isolations at multiple times during the first 100 days after transplant (69 samples and 138 DNA isolations total). These patients represent a sick population since they all received radiation and/or chemotherapy before transplantation, which generally cause organ and tissue damage, especially in the GI tract. Two of the three HCT patients (P07 and P12) also developed graft-versus-host disease (GVHD) in the GI tract and experienced moderate to severe diarrhoea during periods of hospitalisation. Thus collectively across the stool specimens, we were able to cover almost the entire range of the Bristol Stool Scale (types 2-7) with respect to stool consistency.

The median recovery of total DNA (i.e., including both microbial and host DNA) per ~200 μl stool homogenate input was 20 ng with a wide distribution (Fig. 7a). P07 and P12, the subjects who developed GI GVHD, generally showed lower values for total stool DNA than P13, who did not develop GI GVHD (Fig. 7a). We were able to detect both LINE-1 and mt DNA in all stool samples from all three patients. Both the LINE-1 and mtDNA levels in the HCT patients (Fig. 7b-c) showed wider day-to-day variation compared to the healthy individuals (Fig. 6c-f).

**Figure 7.**
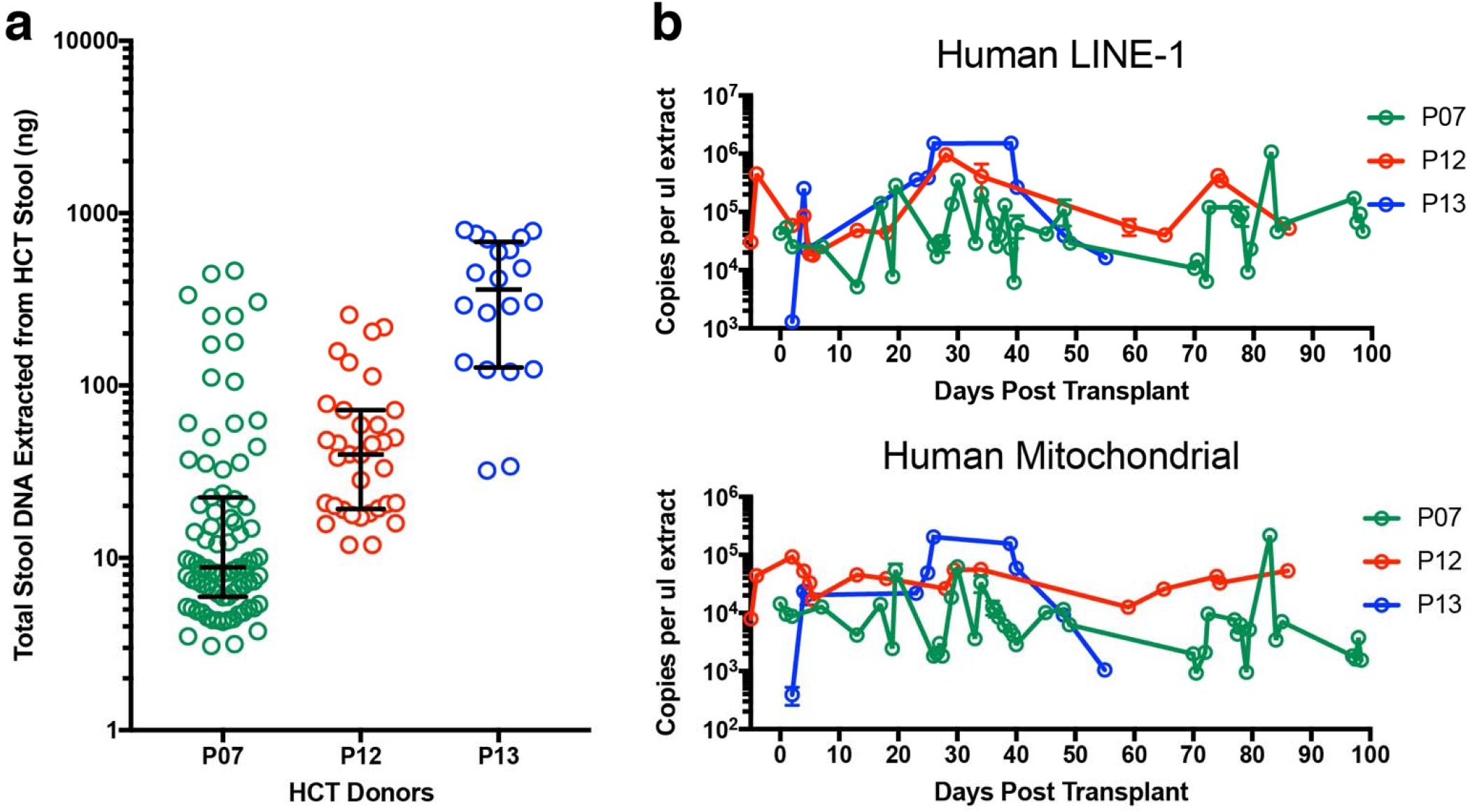
Quantification of host DNA from human patient stools using ddPCR. (a) Total DNA recovered from two side-by-side DNA isolations from 69 stool samples from three hospitalised patients (P07, P12, and P13) undergoing allogeneic HCT. Each circle represent DNA recovered from a single DNA extraction. The horizontal lines represent the median and interquartile range. (b-c) Human DNA quantified using human LINE-1 (60-bp amplicon) and mitochondrial (83-bp amplicon in the ND5 gene) assays. Error bars in (c-f) represent range of copy numbers from single ddPCR reactions on the duplicate DNA extraction. For some points, the error bars would be shorter than the height of the symbol. In these cases, the error bars were not drawn.

For patient P07, for whom Bristol scores were available for every stool, we compared Bristol score to total DNA yield (**Supplementary Fig. S4a**) as well as to ddPCR-assayed host (**Supplementary Fig. S4b**) and 16S (**Supplementary Fig. S4a**) bacterial DNA. The amount of total extracted DNA showed a sustained drop at Day 17 post HCT and coincided with an increase of the Bristol score to 7 (“entirely liquid”). This corresponded to a sharp sustained drop in 16S bacterial DNA as well. In contrast, host DNA levels were maintained throughout the time course, indicating that host DNA can be recovered from liquid stools at comparable levels as seen from low Bristol score stools (**Supplementary Fig. S4b**).

## Discussion

The gut microbiome is implicated in the maintenance of proper health^30^ as well as in nearly every major disease^31^ from obesity, intestinal disorders, to Parkinson’s^32,33^. The host, as a symbiotic partner, interacts intimately with and shapes the microbiota through multiple mechanisms ^34,35^. Yet the lack of well-characterised methods for accessing and analyzing faecal host DNA has limited study of host intestinal effects on the microbiome. It has also limited progress in using stool-based approaches for the study of GI cancer and other diseases affecting the intestinal tract, for example by analysis of methylated DNA that may be recovered in stool.

Here we have developed a pipeline of methods to collect and isolate DNA in stool, and quantify host DNA within stool samples using ddPCR. For sample collection, we identified 0.5 M EDTA (pH 8) for use as a host DNA preservative solution for stool samples, which can stabilize DNA in stool for at least 4 days at room temperature. Since EDTA is nontoxic, readily available and relatively inexpensive, it offers an economical solution for stool DNA preservation at the point of collection, until DNA isolation can be carried out. It is worth noting that our DNA stability analyses were carried out using stool that had been homogenized within an hour of collection, in order to produce material that could be uniformly sampled over multiple time points. In real-world practice, we expect that stools would be collected in EDTA without prompt homogenization. Thus a limitation of our study is that we do not know whether such delays in homogenization would impact the DNA stabilisation effect of EDTA. Additionally, we found glass beads facilitated homogenisation of stool in a relative large volume of solution (i.e. 40 ml) and therefore recommend having them in the stool collectors.

For DNA isolation, we determined that Norgen Stool DNA isolation reagents provided the highest efficiency, nonsize-biased recovery of DNA among the methods we evaluated. For host DNA quantification, we developed four ddPCR assays for quantification of host nuclear and mitochondrial genes in human and mouse stools. The choice of ddPCR as an analytic approach has advantages over real time PCR in this setting. These include obtaining absolute quantification without a standard curve, higher precision^13^, and less sensitivity to PCR inhibitors^36^, which may be present in stool and co-purify with stool DNA^12^. In addition, we chose targets that are present in high copy numbers per cell, and validated low cross-reactivity against other genomes that may be expected in stool. As a result, we achieved high sensitivity (lower detection limit well below a single human nuclear genome), reproducibility, linearity, and specificity with our assays. Ideally, DNA samples should be fragmented into shorter pieces for high CN target analysis (e.g. LINE-1 elements) using ddPCR to avoid target overcrowding in the droplets. However due to low DNA concentration in our patient specimens, we did not perform DNA fragmentation, as incorporating fragmentation may lead to sample loss and/or dilution. Therefore we expect the detection limit to be even lower for the LINE-1 assay for samples that have higher DNA concentrations and are hence suitable for pre-ddPCR DNA fragmentation. When reporting faecal host DNA levels, we found that normalisation of ACN to stool input (wet weight) did not visibly alter the longitudinal trends within an individual, regardless of the individual’s physiological status (healthy vs. hospitalised) and stool consistency (Bristol scores 2 through 7). We infer that this result indicates that the biological variability is much greater than the variability introduced by not normalising to stool weight. However stool wet weight has the limitation that it can be confounded by variations in water content. In future studies, it would be worthwhile to assess whether normalisation to stool dry weight (which was not available for our specimens) could better account for variations in stool input, especially for watery stools that may contain very little organic material.

Using our optimised pipeline, we further demonstrated the feasibility of quantifying host DNA in stools of a wide range of physical properties (as measured by Bristol Stool Scale) from not only healthy populations but also hospitalised HCT patients who commonly have GI tract pathology. In the 31 serially collected stool specimens from three healthy donors, the LINE-1 elements ranged from 5,600,000 to 184,000,000 copies per ~200 μl stool homogenate and 52,000 to 4,050,000 copies per mg of stool collected. Based on the empirically measured number of LINE-1 elements per haploid genome in a commercially available reference human genomic DNA sample, we were able to estimate that these values correspond to approximately 2 to 176 cells per mg of stool collected from the healthy donors. We consider these to be only estimates because the number of copies of LINE-1 per genome is polymorphic in the human population. Thus, there could be differences in the number of LINE-1 copies per genome in the individuals whose stool samples we studied, as compared to the reference human genomic DNA sample. In the healthy individuals studied, the mitochondrial gene ND5 ranged from 1,140,000 to 7,050,000 copies per ~200 μl stool homogenate, corresponding to 9,300 to 182,000 copies per mg of stool collected. In the 69 serially collected stool specimens from the three HCT patients, the LINE-1 elements ranged from 130,000 to 150,600,000 copies (corresponding to approximately 5 to 6544 cells) per ~200 μl stool homogenate. For HCT patients, we could not estimate the number of cells/mg stool because stool weights were not available. The mitochondrial gene ND5 ranged from 39,400 to 21,610,000 copies per ~200 μl stool homogenate in the HCT patient specimens.

Even though many of the stool specimens from HCT patients were watery and had low bacterial counts (see **Supplementary Fig. S4a** for patient P07 as an example), we were able to detect significant amounts of human DNA. We observed day-to-day variations in faecal human DNA in all six human subjects, which were several-fold in healthy controls and up to ~1000-fold in HCT patients. Although technical variation in DNA recovery efficiency does occur among replicate DNA isolations (Fig. 4), this is small relative to the observed day-to-day variations. However in future studies, variation in DNA recovery efficiency could be measured and corrected for using synthetic DNA of artificial sequence spiked into the samples prior to DNA extraction^37^. Taken together, our data suggest that the majority of observed day-to-day variation is likely biological in origin, perhaps related to variation in diet, GI tract environment, GI inflammation or other factors. We speculate that the greater variations seen in the HCT patients may be a manifestation of GI tract damage that is common amongst HCT patients during their treatment course.

Beyond stool, our assays provide a quantitative and reproducible tool that may be applied to quantifying host DNA sequences in other types of human specimens, which may be relevant to a wide range of diseases. Mitochondria are responsible for ATP (energy) generation, reactive oxygen species generation and detoxification, and other essential functions in the cell ^38^. Mitochondrial copy number alterations have been linked to autism ^39,43^ and multiple types of cancer including breast cancer ^41,42^, bladder cancer ^42^, kidney cancer ^42^, and colon cancer ^43,44^. Thus for example, our mitochondrial DNA quantification assays could be adapted to measure such mt copy number alterations in biofluids (e.g., urine, blood, cyst fluid, and cerebrospinal fluid) in cancer patients as a biomarker approach. In addition, our methods could be adapted to other DNA targets, such as specific mutated or methylated DNA sequences in the nuclear genome, for example, which may be associated with cancer and other diseases of the GI tract.

Lastly, as proof-of-concept, we validated that the same approaches for collecting and quantifying human DNA from stool are also effective for quantifying mouse DNA from murine stool. We detected between 253 and 8655 copies of mouse LINE-1 elements in stools from three healthy mice, further demonstrating the functionality of the pipeline. We envision this will facilitate disease studies in a broad range of mouse models of GI diseases. In our experiments, one to three pellets of healthy mouse stools per sample yielded DNA extracts that had to be diluted over 3000-fold for our ddPCR assay. This high sensitivity indicates that mouse DNA will be quantifiable even in sick animals which might produce liquid stools, for example. Taken together, the pipeline of methods for host DNA preservation and detection in stools we have described provides a convenient and highly sensitive tool for quantifying host cell DNA in the GI tract and can be applied broadly in studies of GI tract physiology and disease monitoring.

## Methods

### Stool Collection Kits and Stool Preservatives

Two types of stool collection devices were used for human stool collection in this study. One consists of Precision^™^ Stool Collectors (Covidien, Dublin, Ireland) that were each preloaded with one of two stabilisation buffers: 40 ml of “TEN2” or 50 ml of “EDTA”. TEN2 was made in-house and is composed of 500 mM Tris, 16 mM EDTA and 10 mM NaCl at pH 9.0. EDTA (0.5 M; pH 8.0) was purchased and used without dilution. For specimen collection in the Precision^™^ Stool Collectors, the donors/patients/caregivers were instructed to fill the provided shovel with stool and close the lid with the shovel attached to the lid of the container. The other stool collection device is the OMNIgene Gut kit (DNA Genotek, Kanata, ON, Canada), which has a tube that contains ~ 2 ml of a proprietary buffer and a large stainless steel bead. The stool collection using the OMNIgene kit was conducted per the manufacturer’s instructions.

### Informed Consent and Institutional Review Board (IRB) Supervision of Human Subjects Research

All human subjects in the study (healthy controls as well as patients) provided informed consent and the study was approved by the University of Michigan IRB and carried out in accordance with the relevant guidelines and regulations.

### Healthy Stool Collection and Processing for DNA Stability Testing at Ambient Temperature

Stools from healthy individuals were collected in a hat that sits on the toilet seat. Stool was scooped into an OMNIgene kit and two Precision™ Stool Collectors that we preloaded with TEN2 or EDTA as well as six, 6 mm solid-glass beads. The specimens were then delivered to the laboratory within an hour of bowel movements.

Stool kits were weighed before and after stool collection and stool weight was recorded. Stools were homogenised into slurries as described below and divided into five aliquots. Aliquots were kept at room temperature for a defined amount of time after homogenisation (0, 4, 24, 72, and 96 hours) before freezing at −80°C.

#### Homogenisation for OMNIgene preserved stool

Collection tubes were vortexed vigorously on a Fisherbrand™ Analog Vortex Mixer (Catalog No. 02-215-365) at setting 9 for 30 seconds.

#### Homogenisation for TEN2 preserved stool

Stool weight to buffer volume ratios were adjusted to 1:4 by adding additional TEN2 buffer if stool weight was >10 g. Buffer was not added if the stool weight was <10 g. Stools were homogenised in Precision™ Stool Collectors by vortexing on a Fisherbrand™ Analog Vortex Mixer (Catalog No. 02-215-365) on setting 9 until the sample appeared homogenous to visual inspection.

#### Homogenisation of EDTA preserved stool

Stool weight to buffer volume ratios were unadjusted. Stools were homogenised in the Precision™ Stool Collectors by vortexing on a Fisherbrand™ Analog Vortex Mixer (Catalog No. 02-215-365) on setting 9 until the sample appeared homogenous to visual inspection.

### Healthy Stool Collection and Processing for Longitudinal Human DNA Quantification

The stool was scooped into a Precision™ Stool Collector containing EDTA and six, 6 mm solid-glass beads. Participants were instructed to provide stool samples three times a week. The specimens were then delivered to the laboratory within one hour of bowel movement.

All stool collection kits were weighed before and after stool collection to obtain stool weight. Stools were homogenised into slurries as described below and stored at −80 °C until further analysis.

#### Homogenisation for EDTA preserved stool

Stool weight to buffer volume ratios were unadjusted. Stools were homogenised in the Precision™ Stool Collector by vortexing on high until the sample appeared homogenous to visual inspection.

### Clinical Stool Collection and Processing for Longitudinal Human DNA Quantification

Stools from allogeneic hematopoietic cell transplantation (HCT) patients were collected at multiple time points during the patient’s first 100 days post-transplant, both during hospitalisation by nurses and at home following discharge, in a hat that sits on the toilet seat. Caregivers/patients were instructed to collect bowel movements with a maximum of two per day and to collect a sample of ‘native’ stool for Bristol scoring, as well as to scoop stool into Precision™ Stool Collectors preloaded with 50 ml EDTA without glass beads for subsequent DNA analysis. The specimens were then delivered to the laboratory within 48 hours of bowel movement, at which time the native sample was assessed for Bristol score and the EDTA-stabilised sample was further processed as follows: Stool weight to buffer volume ratios were not adjusted. Stools were homogenised in the Precision™ Stool Collectors by vortexing on a Fisherbrand™ Analog Vortex Mixer (Catalog No. 02-215-365) on setting 9 until the slurry appeared homogenous to visual inspection. The stool slurries were aliquoted and frozen at −80 °C.

### DNA Ladder Extraction and TapeStation Analysis

Invitrogen’s 1kb Plus DNA ladder (100 bp - 15,000 bp) was diluted in TET buffer (10 mM Tris-HCl, 0.1 mM EDTA, 0.05% Tween 20) to concentrations of 50 and 10 ng/μl. A total of 200 μl of each ladder concentration (corresponding to 10 and 2 μg total DNA input) was purified using the Qiagen QIAamp DNA Stool Mini Kit (human DNA protocol), the Zymo Quick-DNA faecal/Soil Microbe Miniprep Kit, or the Norgen Stool DNA Isolation Kit and the ladder eluted in 200 μl elution buffer. One μl of each of the before-purification DNA ladder (corresponding to 50 or 10 ng) and 1 μl of after-purification DNA ladders were loaded onto a High Sensitivity D1000 ScreenTape side by side and analyzed on an Agilent 2200 TapeStation (Santa Clara, CA), such that if DNA recovery were 100%, the same amount of DNA would have been loaded for both the before- and the after-purification samples.

### Oligonucleotide Primers for ddPCR

Primers for the ddPCR analyses of human, mouse, and bacterial DNA targets are provided below in Table 1. Primers for the bacterial 16S ribosomal RNA genes were taken from a published study by Suzuki *et al.^45^*. Primers for human and mouse targets were designed for this study using Primer3 ^46^. Primers were purchased as standard desalted oligonucleotides from IDT (Coralville, IA).

**Table 1.**
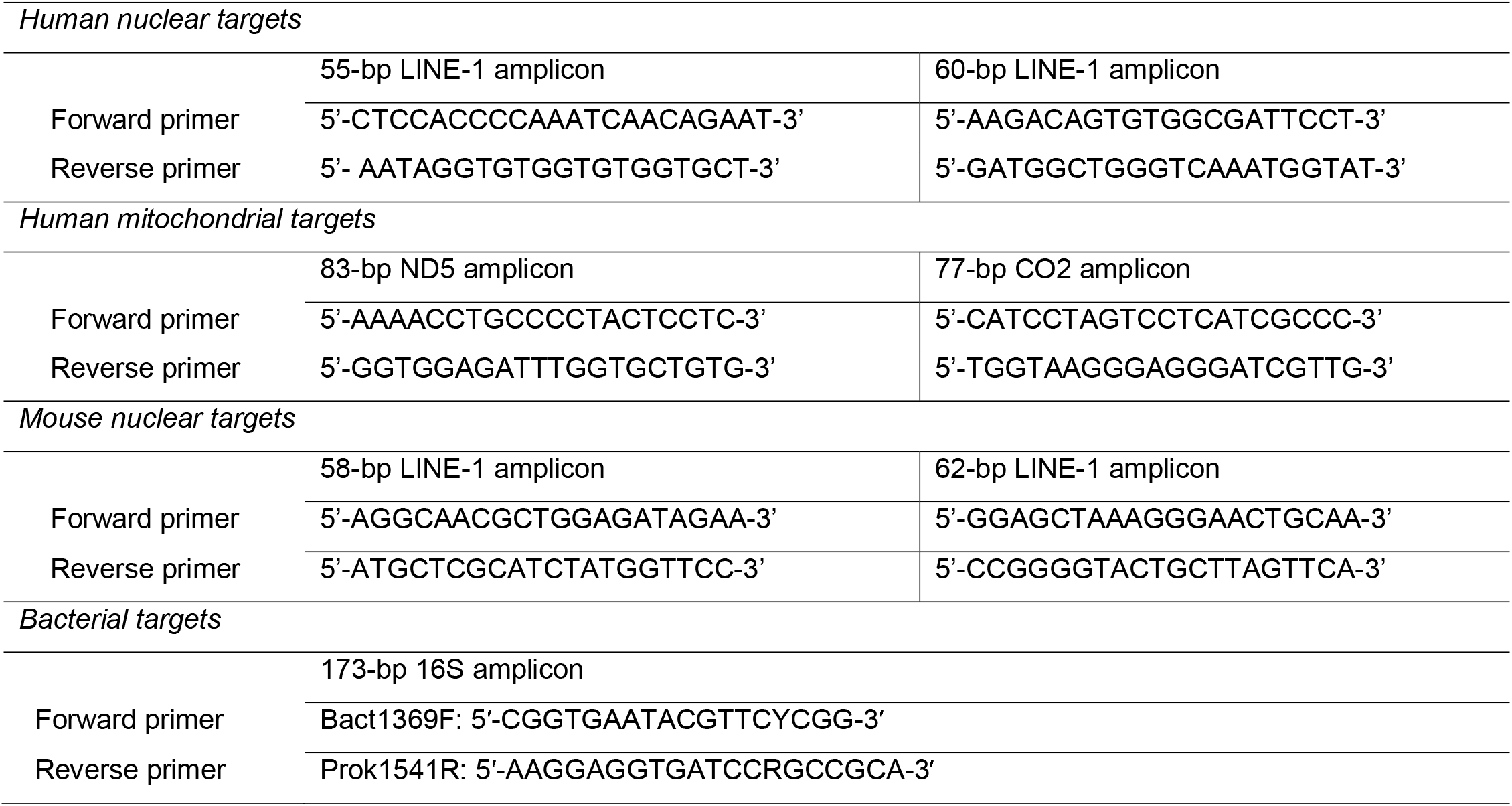
Primer sets used in this study.

### Purified Genomic DNA and Synthetic DNA Fragments

For ddPCR assay development and validation, we used both natural and synthetic templates. The natural templates include purified human (Promega), rhesus monkey (BioChain), mouse (BioChain), corn (BioChain), potato (BioChain), rice (BioChain), bovine (BioChain), *E. coli* (Affymetrix), chicken (BioChain), and wheat (BioChain) genomic DNA (gDNA). Purified human gDNA was digested with the restriction enzyme HaeIII (NEB, Ipswich, Massachusetts) per the manufacturer’s instructions to yield an average of 347-bp DNA fragments^22^. Digestion reactions were frozen at −20 °C until further use without heat deactivation.

The following synthetic gBlock DNA fragments were used for testing linearity, accuracy, and reproducibility for each ddPCR assay:

1. A 126-bp long gBlock (IDT, Coralville, IA) containing the 60-bp LINE-1 amplicon with two modified nucleotides (denoted with underlining) to yield an internal EcoRI site (primer binding sequences are shown in bold):

> 5’-
> ACTGTGAACTAGTTCAACCATTGTGG**AAGACAGTGTGGCGATTCCT**GAGGGATCTAGAA<u>T</u>T<u>C</u>GAA**ATACCATTTGACCCAGCCATC**CCATTACTGGGTATATACCCAAAGGATTATAAATCATGC-3′
2. A 196-bp gBlock (IDT, Coralville, IA) containing the 83-bp ND5 amplicon with two modified nucleotides (denoted with underlining) to yield an internal EcoRI site (primer binding sequences are shown in bold):

> 5’-TCTAGGCCTTCTTACGAGCC**AAAACCTGCCCCTACTCCTC**CTAGACCTAACCTGACTAGAAAAGCTATTACCTAAAA<u>G</u>AATT<u>C</u>**CACAGCACCAAATCTCCACC**TCCATCATCACCTCAACCCAAAAAGGCATAATTAAACTTTACTTCCTCTCTTTCTTCTTCCCACTCATCCTAACCCTACTCCTAATCACATAA-3′

Both gDNA and synthetic gBlocks were diluted serially in TET buffer for ddPCR.

### Stool DNA Extraction for Human Specimens

Total stool DNA was extracted in duplicate from ~200 μl of stool homogenate from each sample using the Stool DNA Isolation Kit by Norgen Biotek (Thorold, Ontario, Canada), Qiagen QIAamp DNA Stool Mini Kit (human DNA protocol), or the Zymo Quick-DNA faecal/Soil Microbe Miniprep Kit per the manufacturer’s instructions. The duplicate purifications for each sample were processed at the same time. Stool DNA concentrations were measured using the Qubit dsDNA HS Assay Kit (Thermo Fisher Scientific).

### DNA Recovery Assessment of Purified Human Genomic DNA Using the Norgen Stool DNA Isolation Kit

*HaeIII*-digested human gDNA was spiked into cleared stool lysates, which were generated in the first phase of the Norgen Stool DNA Isolation protocol, following the initial centrifugation. The protocol was modified slightly for this experiment, in that multiple cleared stool lysates were first pooled. Then, different amounts (40 μl of 800, 80, 8, or 0 ng) of *HaelII*-digested gDNA were spiked into 560 μl aliquots of the pooled lysates. The DNA isolation procedure was completed and subsequently ddPCR for LINE-1 was carried out. The measured ACN for LINE-1 in this scenario comprises both that derived from the spiked-in gDNA, as well as LINE-1 present endogenously in the stool sample. In order to estimate recovery specifically of the spiked-in gDNA, we also measured ACN of LINE-1 present in aliquots of the same stool lysates without gDNA spike-in. This value was then subtracted from the measured ACN in the “with gDNA spike-in” sample to yield the ACN of the spiked-in gDNA. We then calculated % recovery efficiency as: [(measured ACN for spiked-in gDNA)/(expected ACN for spiked-in gDNA)] x 100%. The expected ACN for spiked-in gDNA was determined by extrapolation, based on analyzing serial dilutions of *HaeIII*-digested human gDNA directly using ddPCR.

### Droplet Digital PCR for Human Stool Samples

In a Bio-Rad QX200 droplet generator, 20 μl ddPCR reactions containing 10 μl of 2× QX200™ ddPCR™ EvaGreen^®^ Supermix (Bio-Rad), 1 μl of 5-500 fold TET buffer-diluted stool DNA or 1 μl of TET buffer (for no-template controls, ntc), 0.2 μl of 10 μM forward primer, and 0.2 μl of 10 μM reverse primer were partitioned into ~20,000 oil-emulsified droplets. One technical replicate for each of the TET buffer-diluted duplicate DNA extracts and four technical replicates of ntc were run on ddPCR. The droplets were transferred into 96-well plates and PCR performed using 10 minutes at 95 °C, 40 cycles of 30 seconds at 95 °C followed by 60 seconds at 60 °C, then 5 minutes at 4 °C, 5 minutes at 95 °C, and finally an infinite hold at 4 °C. The temperature ramp increment was 2 °C/second for all steps. Plates were subsequently read on a Bio-Rad QX200 droplet reader.

Gradient PCR was carried out using the same conditions as described above except for using a combined annealing/extension step with no specified temperature ramp increments between steps.

### Mouse Strain and Husbandry

BALB/c (H-2^d^) mice were purchased from Charles River Laboratories (Wilmington, MA). Animal care followed protocols reviewed and approved by the Institutional Animal Care & Use Committee of the University of Michigan, based on the University Laboratory Animal Medicine guidelines. Mice were fed PicoLab 5L0D rodent diet (LabDiet), which is referred to as mouse chow hereafter. DNA from mouse chow was obtained by breaking chow into small pieces with a razor blade and carrying it through the Norgen Stool DNA Isolation kit protocol per manufacturer’s instructions.

### Stool Collection and Processing for Longitudinal Mouse DNA Quantification

Mouse stools were collected twice a week for 32 days. Two hundred μl of 0.5 M EDTA at pH 8.0 was added to each bead tube (part of the Norgen Stool DNA Isolation Kit) and weighed. Each time, one to three pellets of mouse stool was collected directly from the anus of each mouse into the bead tubes containing EDTA, and the tubes weighed again. Stool weight from each collection was calculated for subsequent data normalisation. Within an hour of stool collections, the bead tubes were vortexed to aid in homogenisation and stored at −80 °C until DNA isolation was performed using the Norgen Stool DNA Isolation Kit.

### Stool DNA Extraction for Mouse Specimens

Total stool DNA was extracted once from ~200 μl of stool homogenate from each sample using the Stool DNA Isolation Kit by Norgen Biotek (Thorold, Ontario, Canada) per the manufacturer’s instructions.

### Droplet Digital PCR for Mouse Stool Samples

PCR reactions and droplets were prepared as described for human samples. One μl of 3,600-fold TET buffer-diluted mouse stool DNA and 1 μl of TET buffer were used as templates and NTCs (no template controls), respectively, in the PCR reactions. PCR was carried out as follows: 10 minutes at 95 °C, 40 cycles of 30 seconds at 95 °C followed by 60 seconds at 59 °C, then 5 minutes at 4 °C, 5 minutes at 95 °C, and finally an infinite hold at 4 °C. The temperature ramp increment was 2 °C/second for all steps. Plates were subsequently read on a Bio-Rad QX200 droplet reader.

### Droplet Digital PCR Data Analysis

QuantaSoft (Version 1.7.4.0917) was used for raw data processing. All samples included in analysis had a minimum of 10,000 accepted droplets during the ddPCR reading. The thresholds for positive and negative droplets were set as follows: 3400 for the four human target assays (the 55-bp and 60-bp human LINE-1 assays, and the 77-bp and 83-bp human mt assays), 6300 for the 172-bp bacterial 16S assay, and 3000 for the two mouse target assays (the 62-bp and 58-bp mouse LINE-1 assays). These thresholds, although chosen arbitrarily, were set using droplet fluorescence values from NTC samples to define what would be considered negative droplets. The copy numbers per reaction of template-containing samples were averaged between the two extraction-duplicates to yield ACN_samp_ı_e_. The copy numbers per reaction of ntc were averaged among the four technical replicates to yield ACNntc. Then copy numbers of the template-containing samples were adjusted for background and dilution using the equation: (ACN_sample_ - ACN_ntc_)^*^dilution-factor, which give the copies per μl DNA extract. To normalise to stool input, copies per μl DNA extract were divided by the mass of stool that was contained in the ~200 μl stool homogenate being added to each extraction.

## Acknowledgements

We thank Elena Stoffel for assistance with IRB approvals; Missy Tuck, Kirk Herman, and Annika Goicochea for coordinating specimen collections; David Hyland, Ryan Lindstrom, Tyler Suciu, Joshua Weber-Townsend, Ricardo Engel, and Ruta Raulickis for assistance with specimen processing; and Nicholas W. Lukacs and Judith Luborsky for critical comments and assistance in editing the manuscript.

K.H. was supported by a Clinical and Translational Science Award (CTSA) from the National Institutes of Health. H.F. was supported by JSPS Postdoctoral Fellowships for Research Abroad and The YASUDA Medical Foundation Grants for Research Abroad. M.T. and S.W.C. acknowledge support from an A. Alfred Taubman Medical Research Institute Grand Challenge Award. P.R. acknowledges support from National Institutes of Health grants CA203542; CA217156; HL128046. M.T. acknowledges support from Mary Petrovich through the University of Michigan Fast Forward Gastrointestinal (GI) Innovation Fund and from the University of Michigan Comprehensive Cancer Center Support Grant.

## Author contributions statement

K.H., P.R., S.W.C., and M.T. conceived the study and interpreted results. K.H., E.S., and M.T. designed the experiments. K.H., H.F., and C.Z. performed the experiments. K.H. and M.T. analysed the results. K.H., E.S., and M.T. wrote the manuscript. All authors reviewed the manuscript.

## Data availability

The data generated or analysed during this study are included in this published article in the Figures and Supplementary information files.

