## Supplementary info for "A Pipeline for Faecal Host DNA Analysis by Absolute Quantification of LINE-1 and Mitochondrial Genomic Elements Using ddPCR"

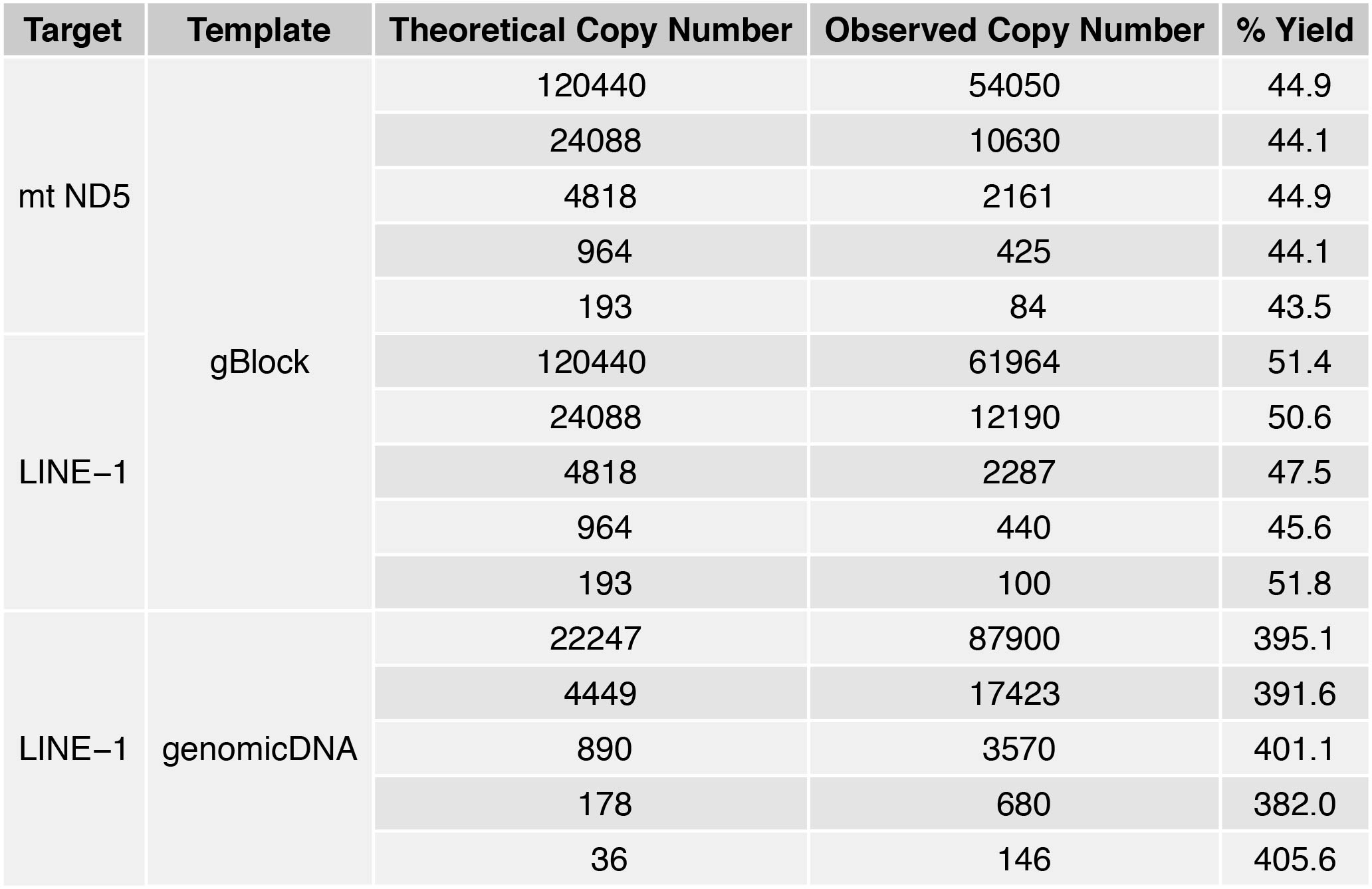
**Supplementary Table S1**. Expected copies based on known amounts of input (i.e., “Theoretical Copy Number”) as well as Observed Copy Number and % Yield for each of the points on the graphs in **Fig. 2a-c**.

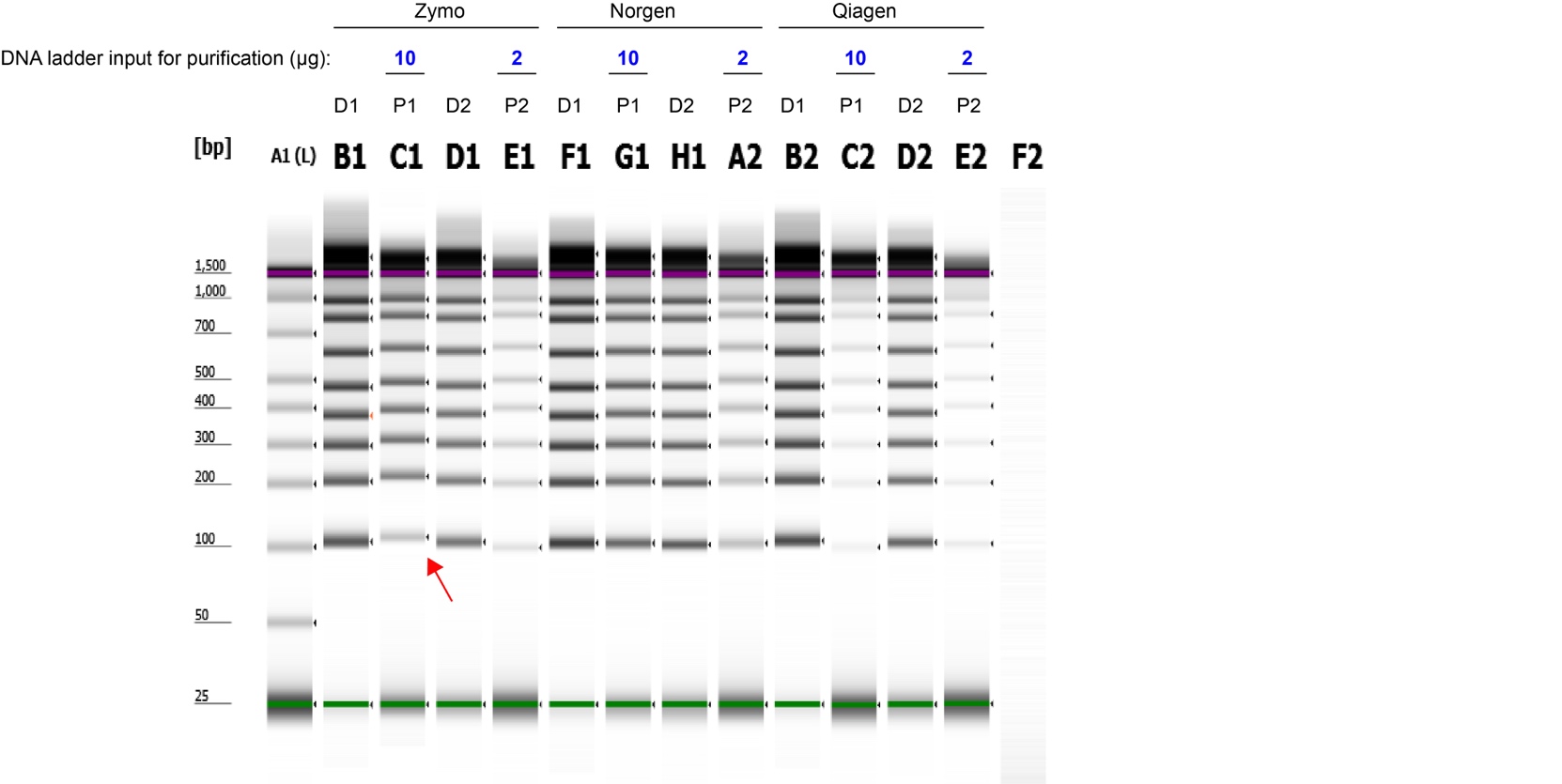

**Supplementary Figure S1.** The uncropped, full-length gel image shown in **Figure 3b**. For direct visualization of DNA recovery from the three different purification processes, DNA ladder was either loaded directly onto TapeStation (D) or was subjected to one of the three purification protocols before being loaded onto TapeStation (P). In a 100% recovery scenario, the D and P lanes should contain the same amount of DNA. The purple and green lines in each lane represent internal upper and lower markers, respectively, for sizing and alignment. The red arrow denotes low recovery of the 100 bp band from the Zymo-extracted ladder samples.

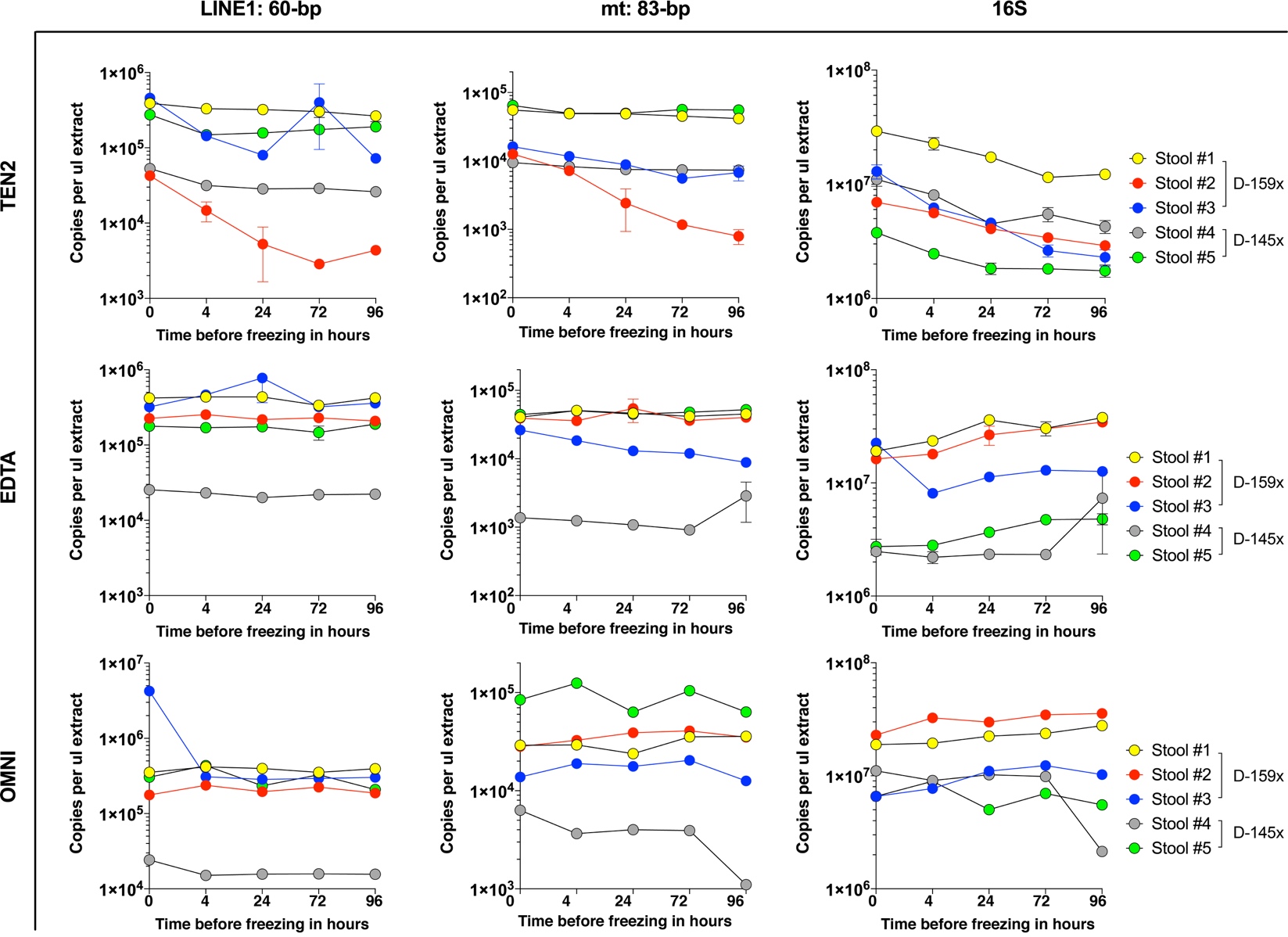

**Supplementary Figure S2.** Effectiveness of different stool preservation solutions on endogenous DNA stability. Absolute copy numbers of LINE-1 (left), mt (middle), and 16S (right) DNA are plotted for varying incubation times of stool specimens at room temperature.

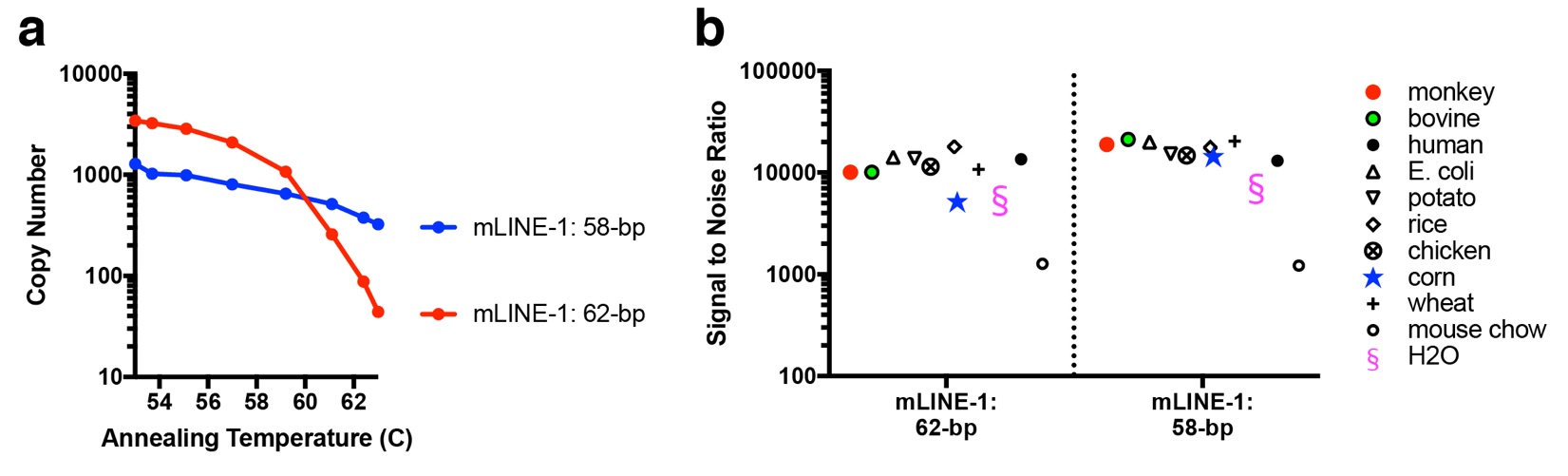
**Supplementary Figure S3**. Optimisation of PCR assay conditions and specificity assessment for the faecal mouse DNA ddPCR assays. (a) Annealing temperature optimisation and (b) DNA specificity evaluation, using two sets of primers targeting mouse LINE-1 elements.

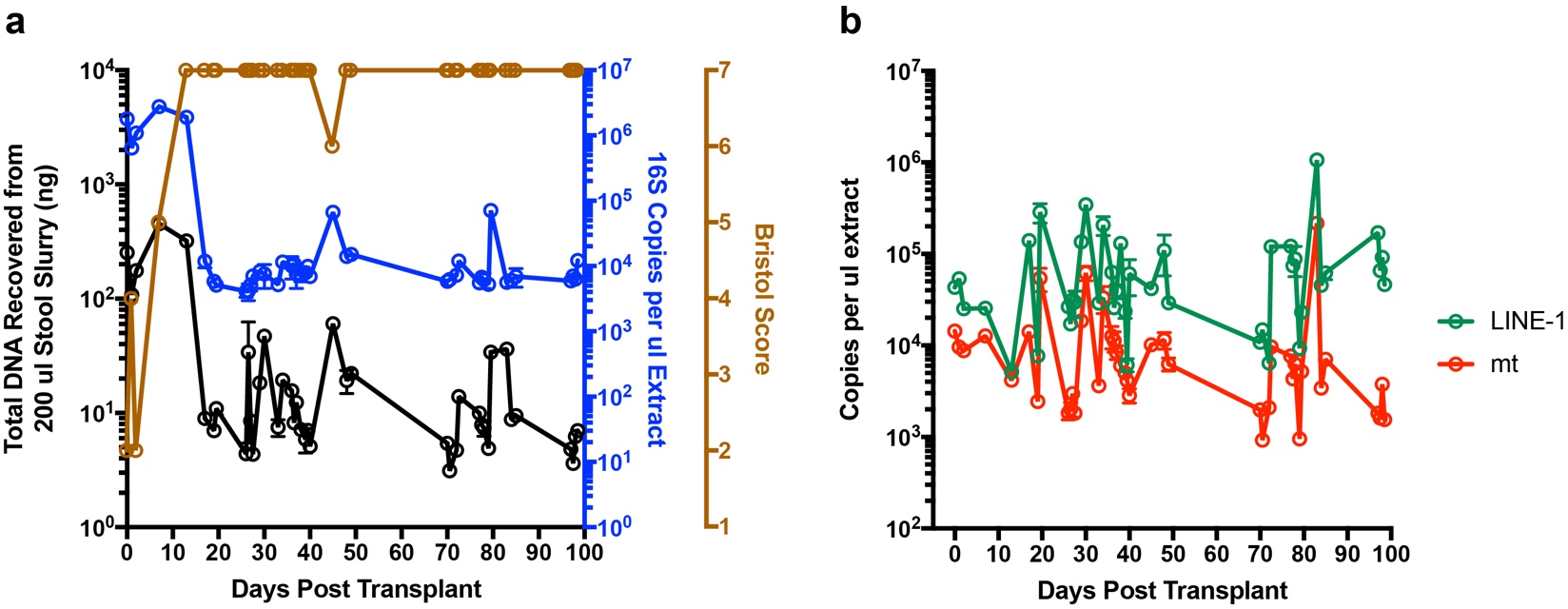
**Supplementary Figure S4**. Characterisation of longitudinal stool specimens collected from a hospitalised patient (P07) who had undergone HCT. (a) Total DNA recovered from each stool slurry (in black), ddPCR quantification of 16S bacterial genes in each stool DNA extract (in blue), and Bristol scores for each stool specimen (in brown). (b) ddPCR quantification of LINE-1 (in green) and mtDNA (in red) in each stool DNA extract. DNA was extracted twice side-by-side from each stool sample, and the DNA extracts were each analyzed once on ddPCR. The error bars represent the range of DNA recovered (in black in **a**) or copies detected (in green and red in **b**) from the duplicate extracts for each stool. The Bristol scores do not have error bars as they were assessed once for each specimen. For some points, the error bars would be shorter than the height of the symbol. In these cases, the error bars were not drawn.
